# Single Objective Light Sheet Microscopy allows high-resolution *in vivo* brain imaging of Drosophila

**DOI:** 10.1101/2024.11.06.622263

**Authors:** Francisco J. Tassara, Mariano Barella, Lourdes Simó, M. Mailén Folgueira Serrao, Micaela Rodríguez-Caron, Juan Ignacio Ispizua, Mark H. Ellisman, Horacio O. de la Iglesia, M. Fernanda Ceriani, Julián Gargiulo

## Abstract

*In vivo* imaging of dynamic sub-cellular brain structures in *Drosophila melanogaster* is key to understanding several phenomena in neuroscience. However, its implementation has been hindered by a trade-off between spatial resolution, speed, photobleaching, phototoxicity, and setup complexity required to access the specific target regions of the small brain of *Drosophila*. Here, we present a single objective light-sheet microscope, customized for *in vivo* imaging of adult flies and optimized for maximum resolution. With it, we imaged the axonal projections of small lateral ventral neurons (known as s-LNvs) in intact adult flies. We imaged the plasma membrane, mitochondria, and dense-core vesicles with high spatial resolution up to 370 nm, ten times lower photobleaching than confocal microscopy, lower invasiveness and complexity in sample mounting than alternative light-sheet technologies, and without relying on phototoxic pulsed infrared lasers. This unique set of features paves the way for new long-term, dynamic studies in the brains of living flies.

## Introduction

*Drosophila melanogaster* is an ideal model to address key questions in neuroscience, given its relatively simple and genetically tractable nervous system. The recent availability of the fly brain connectome predicts new lines of research requiring live brain imaging of intact individuals.^1^ Recent advances in optical microscopies have enabled impressive *in vivo* mapping of neuronal activity maps, and their correlation with sensory stimuli, behaviour and metabolism.^2,3^ In addition, the community has built expertise on how to create fly preparations for long-term optical brain imaging^4–6^ and the development of protein-based probes.^7^ Despite this, long-term imaging of dynamic sub-cellular anatomical structures such as individual neurites, cytoskeleton or organelles remains less explored. This approach is key to understand phenomena such as circadian structural plasticity,^8,9^ time of day-dependent connectivity,^10,11^ axonal regeneration and lesion recovery,^12^ or the structural correlates that result from forced depolarization/hyperpolarization. However, achieving long-term structural imaging requires the development of novel methods offering high spatial resolution, good optical sectioning, minimal photobleaching and phototoxicity, and low invasiveness.

Currently, the most popular approach to *in vivo* brain imaging in *Drosophila* is two-photon microscopy (2pM).^13–15^ With 2pM, it has been possible to acquire brain activity maps that are odor-^2,5,16^ auditory-,^17^ or visually-evoked,^18,19^ during walking,^20,21^ sleeping,^22^ or eating.^23^ In addition, it allowed the study of functional correlations between brain regions,^3^ and their correspondences with metabolism.^24^ However, 2pM has significant limitations for imaging living animals. Firstly, the use of high peak pulse powers can induce photodamage or brain heating,^25^ or even behavioral responses.^20^ Secondly, it induces substantial photobleaching that limits the number of volumes that can be acquired. Pulsed lasers can be avoided using one-photon microscopies such as confocal,^26^ spinning disk^27,28^ or epifluorescence with light-field.^29–31^ However, these techniques can suffer from similar or even higher photobleaching, or higher optical and computational complexity.

The technology with the potential to alleviate most of the aforementioned drawbacks is light sheet (LS) fluorescence microscopy,^32–35^ due to its high optical sectioning and low phototoxicity compared to scanning methods. However, there are several challenges to implement this technique for *in vivo* high-resolution brain imaging in adult *Drosophila*. Conventional LS microscopes require two perpendicular objectives, which complicates the interface with the head of the fly, restricts the imaging numerical aperture (NA), and complicates sample scanning. For these reasons, the implementation of LSM to adult *Drosophila* has been limited to very low spatial resolution^36^ or *ex vivo* preparations.^37^ Very recently, a new family of LS microscopes has emerged that use the same objective for illumination and detection. They are known as Single Objective LS (SOLS)^38–43^ or Swept Confocally-Aligned Planar Excitation (SCAPE).^44,45^ SCAPE microscopy leveraged the implementation of LSM to living adult animals such as mice,^44^ *C. elegans*,^45^ and zebrafish.^45^ Very recently, SCAPE demonstrated brain-wide calcium imaging of living *Drosophila* adults at 10 Hz.^46^ However, this research was focused on mapping the activity of clusters of neurons ranging from a few to tens of microns in size, with suboptimal spatial resolution and an imaging voxel size of (1×1.4×2.4) μm^3^.

Here, we designed and built a SOLS microscope tailored for *in vivo* imaging of adult *Drosophila* and optimized for maximum spatial resolution. It allows brain imaging with a high NA objective and no mobile parts at the interface with the fly. With it, we demonstrated *in vivo* imaging of the axonal projections of small lateral ventral neurons (s-LNvs), with subcellular high spatial resolution up to 370 nm. We imaged the plasma membrane, mitochondria and dense-core vesicles (DCVs). In addition, we tested the suitability of the technique for long-term imaging. We demonstrate that under typical imaging conditions, the fluorescence emission drops to 50% after the acquisition of 80 3D volumes. Furthermore, we present imaging during time lapses of up to two hours. Overall, the built microscope offers high spatial resolution in combination with the convenient sample mounting of confocal or 2p microscopy and the low photobleaching and phototoxicity of LS illumination, making it ideal for long-term structural studies of living flies.

## Results

### Experimental setup

Figure 1a shows the schematics of the SOLS microscope with oblique illumination, optimized to facilitate the interface with the living fly, and to maximize spatial resolution. In short, it consists of an upright primary microscope with a water immersion objective O1 (60×, 1.1 NA) that excites the sample with oblique light-sheet illumination and collects the fluorescence emission, which is relayed by a remote focusing system that creates a virtual replica of the sample using secondary objective O2 (60×, 0.9 NA, air). A remote microscope equipped with a bespoke glass-tipped objective O3 (1.0 NA) is oriented at an angle and images the virtual replica.^41^ All microscope modules are configured as 4f optical systems. The experimental setup also features an epifluorescence wide-field (WF) microscopy module, useful for coarse sample positioning. A flip mirror (M’) and a beam splitter (BS) can be placed on the excitation and detection paths to switch between the LS and WF configurations.

**Figure 1:**
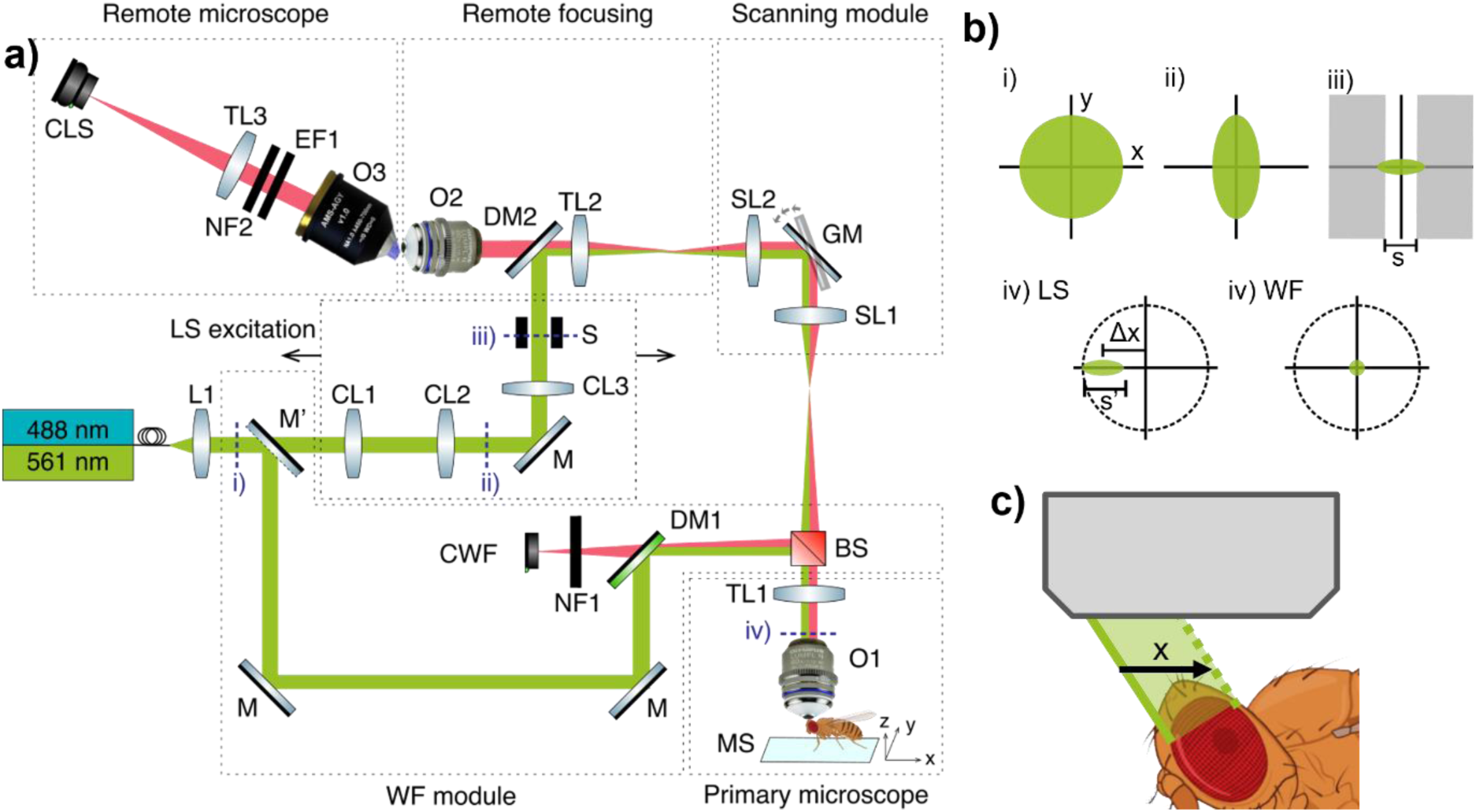
**a)** Schematic of SOLS. The experimental setup is divided into main key modules: The LS excitation module shapes the laser beam for oblique plane illumination; the scanning module is responsible for sweeping the sheet of light across the sample; the primary microscope illuminates the sample and detects the fluorescence signal; the remote focusing system creates a virtual replica of the sample; the remote microscope images the replica. The microscope can be set in an epifluorescence WF configuration (WF module) for coarse sample positioning. O1, O2 and O3: primary, secondary and tertiary objectives, respectively. TL1, TL2, TL3: Tube lenses. DM, DM2: Dichroic mirrors. CL1, CL2, CL3: Cylindrical lenses. M: mirror. M’: Flip mirror. GM: Galvanometric mirror. BS: Beam splitter. SL1, SL2: Scanning lenses. CWF: Camera used in Epifluorescence mode. CLS: Camera used in Light Sheet mode. EF1, EF2: Emission filters. NF: Notch filter. MS: Motorized stage. Arrows in the LS excitation module depict that the module is built on top of a translation stage. **b)** Sketch of the beam shape at different positions along the optical path. Panels i to iv are not to scale and correspond to the planes indicated with blue dashed lines in (a). Panel iv illustrates the beam shape for LS (left) and WF (right) illumination. The real sizes of the beam at each panel, calculated as the Full Width Half Maximum in (x, y) are: i) (2.35, 2.35) mm, ii) (0.47, 2.35) mm, iii) (0.47 mm, focused). Slit width s=0.8 mm. iv-WF) (s’=1.06 mm, focused), iv-WF) (focused, focused). **c)** Sketch representing the light sheet scan over the sample space.

To create the LS, the beam is shaped by the LS excitation module. To facilitate the description of the beam shaping and routing, Figure 1b shows the cross-section of the beam at selected planes, marked with dashed blue lines in Figure 1a. The excitation uses a fiber-launch system that combines two continuous-wave, linearly polarized, 488 nm and 561 nm lasers. An achromatic lens L1 (f=30 mm) collimates the beam to a circular Gaussian profile (Figure 1b, panel i). Then, the beam is reduced 5× in the *x*-axis by two cylindrical lenses, CL1 (f=250 mm) and CL2 (f=50 mm), as shown in Figure 1b, panel ii. A third cylindrical lens CL3 (f=100 mm) focuses the beam on the *y*-axis onto an adjustable mechanical slit, as shown in Figure 1b, panel iii. The slit is placed in a plane that is conjugated to the back focal plane of the objective O1 (Figure 1b, panel iv-LS). In this way, the slit truncates the Gaussian beam in the *x*-axis, and its width defines the excitation NA of the LS. The entire LS excitation module is mounted on a translation stage that shifts off-axis the beam along the *x*-axis at the back focal plane of O1 by a magnitude *Δx*. This offset controls the LS angle θ.

In the WF module, the excitation beams pass through a 4× beam expander and are later focused at the back focal plane of O1 (Figure 1b, panel iv-WF), resulting in a field of illumination (FOI) of ∼370 μm.

To achieve non-distorted remote focusing, the composed magnification M1-2 between the sample and its virtual replica must satisfy M_1-2_=*n*_2_⁄*n*_1_, where *n*_1_ and *n*_2_ are the refractive index at the sample and image planes, respectively.^47,48^ This is achieved by using a Tube Lens (TL2=135 mm) that is the assembly of three optical components (for details, see Materials and Methods and Millett-Sikking *et al.* ^49^). Finally, the remote microscope is oriented so that its axis forms an angle θ with respect to the O2 axis. In this way, the imaging plane is orthogonal to the excitation, which is the standard requirement in LS microscopy.

The scanning module uses a single axis-galvanometric mirror (GM) to perform lateral scanning of the LS through the sample in an angle-invariant way, as schematized in Figure 1c and explained in detail by Kumar *et al*.^50^ The GM plane is conjugated to the back focal plane of both objectives O1 and O2. This configuration enables the GM to de-scan the fluorescence emission, keeping the virtual image generated by the remote focusing module stationary. During the scanning of the LS in the *x* direction, an image is acquired with the LS camera (CLS) for each LS position. This creates a 3D parallelepiped-shaped volume (Figure 1c, light green box) that is later computationally deskewed into the traditional cartesian coordinates system.

### Instrument characterization

Figures 2a and 2b show the schematics of the illumination light sheet and its relevant dimensions. It is a truncated Gaussian beam with an incident angle *θ* with respect to the *x*-axis. The beam propagates parallel to the *z’* axis, is collimated in the *yz’* plane and is focused on the *x’z’* plane. It is defined by a beam waist *w0*, a width *wy* and a length L*z’* in the *x’*, *y* and *z’* axis, respectively. In SOLS microscopy — and LS microscopy in general — there is a trade-off between resolution, sectioning, axial FOI, and collection efficiency, which depends on the LS incident angle and NA. In this setup, we have optimized the design to maximize resolution and sectioning at the expense of axial FOV. The angle was set to *θ*=41°, following the criteria of matching the LS thickness to the axial point spread function (PSF) of the emission, as described by Millett-Sikking *et al.*^49^

**Figure 2:**
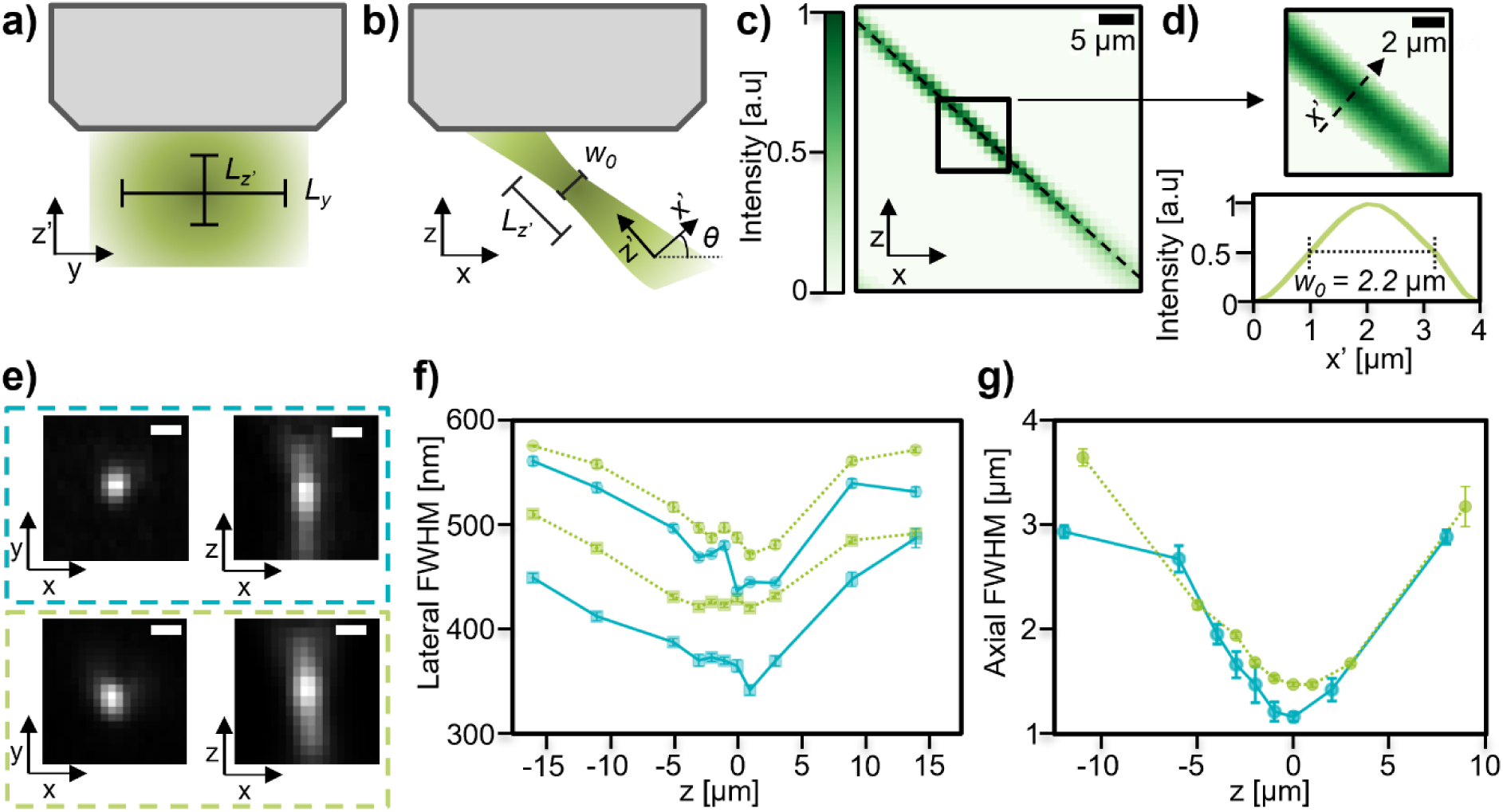
Characterization of the excitation light sheet and the microscope resolution. a) and b) Sketch of relevant dimensions of the LS illumination. c) Normalized intensity of the measured light sheet beam in the xz plane. d) Top: Detail of the light sheet beam waist shown in the black box of (c) Bottom: Normalized intensity profiles of the LS in the x’ axis. e) Top, blue dashed box: PSF cross-section in the xy and xz planes for a 488 nm excitation. Bottom, green dashed box: for a 561 nm excitation. Scale bar:1 μm. f) Lateral PSF FWHM as a function of z for 488 nm (light blue solid line) and 561 nm (light green dashed line). Circular and square scatters represent the PSF width in the x and y directions, respectively. g) Axial PSF FWHM as a function of z for 488 nm in blue and 561 nm in green.

The beam profile was characterized by scanning a single 100 nm fluorescent bead through the LS. For each position of the bead, the total detected fluorescence emission at the LS camera was calculated. Figure 2c illustrates the resulting normalized intensity map, acquired at an excitation wavelength of 561 nm. The measured angle is *θ*= (41 ± 1)°. Figure 2d shows an intensity map of the region closer to the beam waist and a linear profile along the *x*’ axis. The measured full width at half-maximum (FWHM) is *w_0_=* (2.2 ± 0.1) μm. The width of the LS is L_y_*=*(53 ± 1) μm calculated as the FWHM and the length is Lz’= (39 ± 1) μm computed as twice the distance over which the intensity drops by a factor of 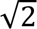 (see Supplementary Note 1 for details). It must be noted that the Field of View (FOV) in the *x* axis is given by the length of the scan, only limited by the FOV of the primary objective (∼441 μm for the 60x objective used here). On the other hand, the FOVy in the *y* axis is determined by the FOV of the remote focusing system (∼113 μm for the presented configuration, see Materials and Methods for details).

We studied the axial and lateral resolution for both excitation lasers. For that, a 2D calibration sample consisting of individual beads drop-casted on a glass substrate was used (see Materials and Methods for details). The sample was placed in a series of *z* positions relative to the focal plane of the objective. For each *z* position, a 3D volume was imaged, and the PSF of each bead was analyzed. Representative PSFs are shown in Figure 2h for both excitation lasers at *z* = 0. The full width at half maximum FWHM was computed for each bead in the three axes. Figure 2i shows the lateral FWHM for both lasers as a function of the z position of the sample. The lateral resolution is relatively uniform in an axial range of approximately 30 μm. The resolution is slightly better for the *x*-axis (solid lines) than for the *y*-axis (dashed lines). This is a typical feature of SOLS microscopy and is explained by a loss of effective detection NA in the direction of the tilt of the remote microscope (see Supplementary Note 3 for further discussion).^41,45^ At the focal plane, the resolution was (FWHM_*x*_, FWHM_*y*_) = (365 ± 6, 437 ± 3) nm, and (FWHM_*x*_, FWHM_*y*_) = (429 ± 3, 488 ± 5) nm for the 488 nm and 561 nm excitation, respectively. Figure 3j shows the axial resolution as a function of the *z* position of the sample. At the focal plane, the FWHM_*z*_ are (1150 ± 40) nm and (1460 ± 30) nm for 488 nm and 561 nm excitation wavelengths, respectively. Away from this plane, the LS increases its thickness, deteriorating the axial resolution. The obtained values are consistent with the Rayleigh length *R_l_* of the LS,^51^ *R_l_* = 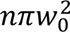⁄(8 λ *ln*2) = (6.5 ± 0.3) μm.

**Figure 3:**
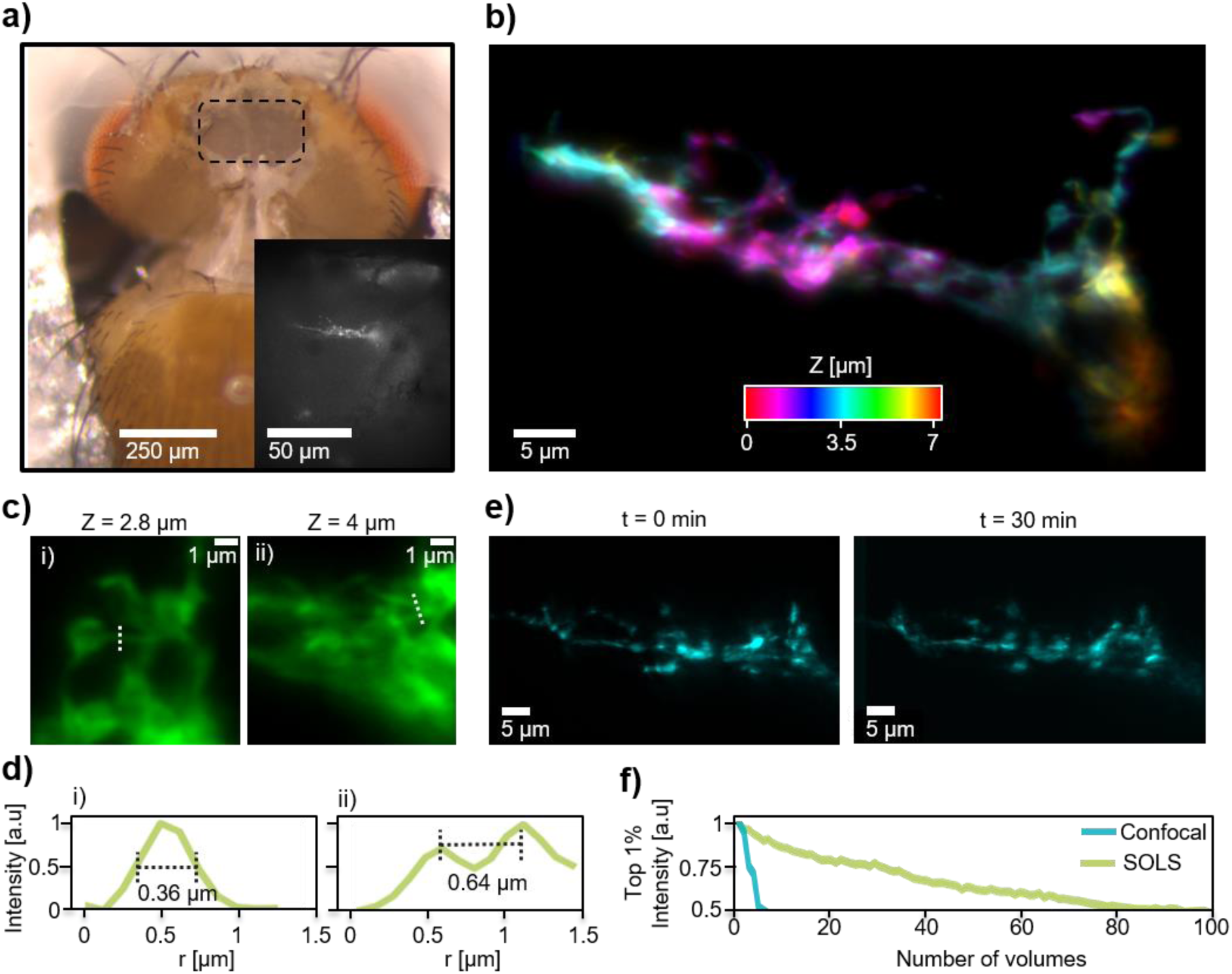
Imaging fluorescent sLNv neurons in living adult flies. **a)** Image showing the mounted fly, highlighting the area where the cuticle dissection was performed with black dashed lines. The inset shows s-LNv terminals imaged with the WF fluorescence module. **b)** Representative image of s-LNv terminals acquired with the LS mode of the microscope. The axial position is color coded. **c)** Magnified views of distinct regions within the terminals at selected axial frames, to reveal small structural details. **d)** Intensity profiles across the white-dashed lines marked in (c)**. e)** s-LNv terminals imaged for 1 hour with a 1-minute interval. The first image is shown in the left, and the one after 30 minutes in the right. Both images are z-projections of the entire volume. **f)** Normalized intensity of the brightest 1% pixels in each volume as a function of the number of imaged volumes for a confocal microscope (blue) and SOLS (green). All images of this figure correspond to flies expressing ;UAS-mCD8::GFP;Pdf-GAL4 excited at 488 nm

### High-resolution and low-bleaching imaging of sLNv neurons in living adult flies

To demonstrate the capabilities of the SOLS microscope in a living fly, we imaged the axonal projections of the s-LNvs. These neurons are a key component of the circadian clock in the *Drosophila* brain as they coordinate circadian locomotor activity in constant conditions.^52^ They also release the neuropeptide pigment-dispersing factor (PDF), which is necessary to maintain the internal coherence of the circadian network.^53^ The somas of the s-LNvs are in the accessory medulla and project to the dorsal protocerebrum, where they contact other clusters of clock neurons.^54^ These projections are very stereotyped and have genetic drivers for their manipulation and specific fluorescent labeling. In addition, their projections are found approximately 25 μm below the brain surface, which makes them accessible during *in vivo* preparations.

For imaging, the flies were mounted on an aluminum foil with a hand-made triangular hole, as shown in Figure 3a. First, each fly was anesthetized on ice and gently manipulated to position the head and thorax into the hole using forceps under a dissection microscope. Then, the head was bent forward to give access to its posterior surface and carefully immobilized by the eyes employing beeswax at 70 °C. Later, a small portion of the cuticle of the dorso-posterior region of the head was removed under saline solutions (mark with a dashed square in Figure 3a, see Methods for details). Finally, the air sacs and the fat tissue were also removed to obtain optical access to the s-LNv region. Importantly, the M16 muscle was cut to reduce brain motion.^55^

We imaged transgenic flies expressing plasma membrane GFP in the s-LNv neurons (;UAS-mCD8::GFP;*Pdf*-GAL4). The inset of Figure 3a shows a representative image of the neurons as seen by the WF mode, useful for positioning the region of interest to be imaged by the LS mode. Then, 3D volumes were acquired at an excitation wavelength of 488 nm, with a FOI of 80×50×18 μm^3^ and a voxel size of 0.12×0.12×0.16 μm^3^. A representative top view (*xy*) of the right terminals of the s-LNv neurons is shown in Figure 3b, with the axial position color-coded. A 3D projection of the data is shown in Supplementary Video 1. To show the resolving power of the microscope, Figure 3c shows close-ups of a few selected frames from the same 3D volume, presenting small features. Figure 3d-i shows a line intensity profile of a selected thin process with a FWHM of 360 nm. Figure 3d-ii shows a line intensity profile across two resolved thin processes, separated by 640 nm. Furthermore, a parameter-free image resolution estimation based on decorrelation analysis was implemented, following the methods from Descloux *et al.*^47–49^. The estimated lateral resolution of the *z*-stack is 540 ± 10 nm (see Supplementary Note 3 for details). The algorithm does not distinguish between the *x* and *y* directions, and therefore, the value corresponds to an average lateral resolution.

We studied the suitability of the technique for long-term imaging. The plasma membrane GFP in the s-LNv neurons of a living fly was imaged for 60 minutes, with a 1-minute interval between them. Figure 3e shows the same area of the sample, at the beginning of imaging (left) and after 30 minutes of imaging (right). Notably, the image is well visible without significant signal loss. Then, to quantitively study the photobleaching of the sample, flies were exposed to 100 consecutive 3D-acquisitions. Figure 3f (green line) shows the intensity of the brightest 1% pixels in the volume, as a function of the number of acquisitions. Typically, the intensity drops to half after (75 ± 10) acquisitions. It must be noted that the bleaching rate depends on the acquisition parameters, such as the integration time or the excitation irradiance. Increasing any of them provides images with higher signal to noise ratio (SNR) at the expense of faster photobleaching. For these experiments the SNR was (26 ± 4). For comparison, an equivalent experiment with living flies was performed using a confocal microscope. The acquisitions parameters were set to obtain images with a similar SNR, which was (22 ± 4). Figure 3f (blue line) shows that in the confocal microscope, intensity drops to half after (6 ± 2) acquisitions. This means that the SOLS microscope presented here allows the acquisition of ten times larger number of volumes than the confocal, with similar image quality. For further details on the photobleaching experiments, see Supplementary Note 4.

We then performed two-color imaging of the s-LNv neurons using the two available excitation lasers. First, we imaged flies expressing GFP in the plasma membrane and mKO fluorescent protein in the mitochondria.^56^ Figure 4a shows an example of a *z*-projection in an axial range of 7 μm. GFP is shown in green and mKO in red. Figure 4b shows a close-up of a single selected frame. On the other hand, most of the mKO signal appears in small clusters with sizes ranging from 400 to 800 nm, which is consistent with the typical size of a mitochondrion at this time of _day._56,57

**Figure 4:**
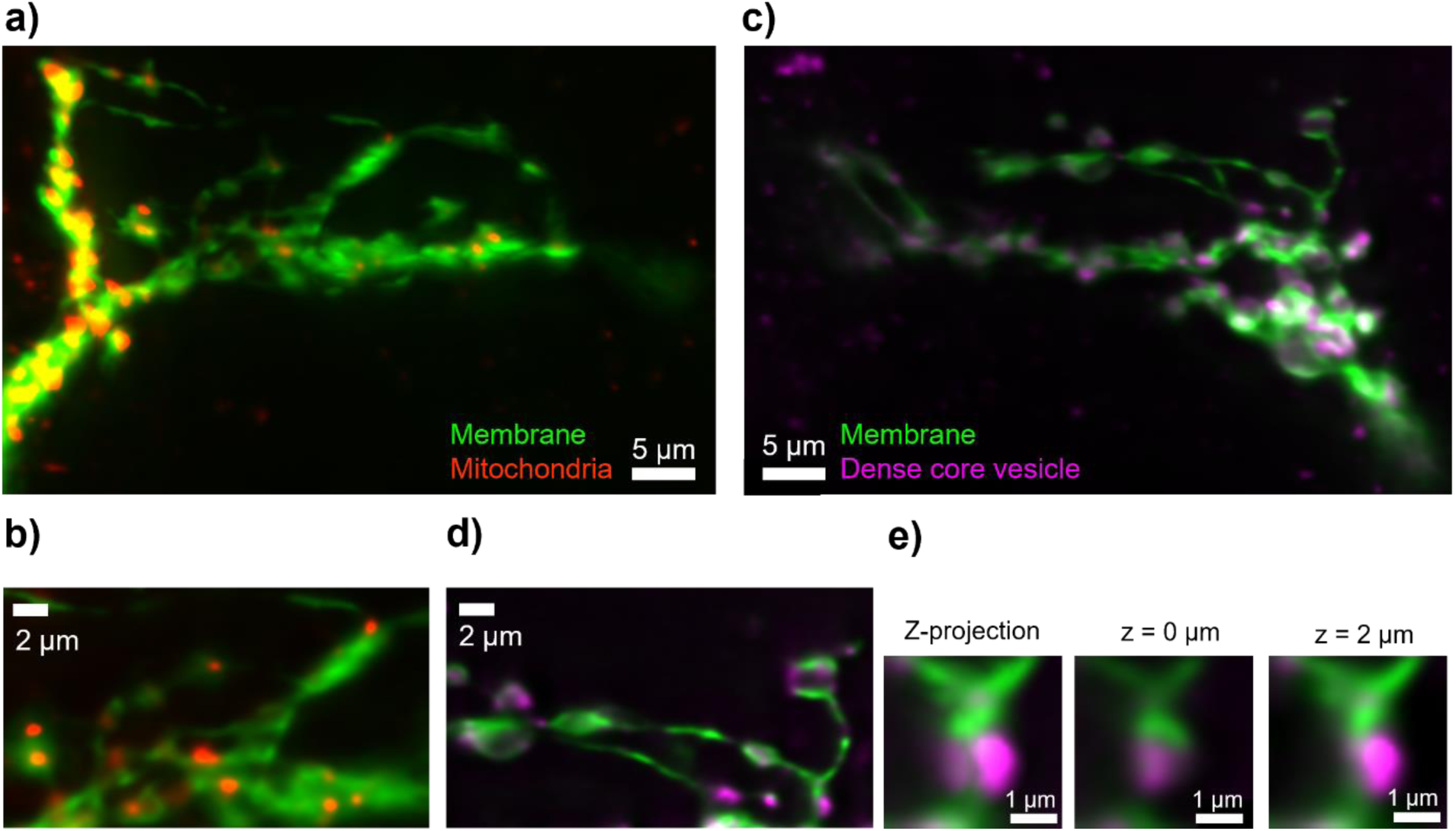
**a**) Image of an sLNv terminal in a fly expressing ;UAS-mCD8::GFP;Pdf-GAL4 > ;UAS-mito::mKO;. The membrane is labeled in green (GFP), while mitochondria are labeled in red (mKO). **b**) Magnified views of a single axial frame at a selected region from panel (a), showcasing finer structural details of the mitochondrial and membrane organization. **c**) Image of an sLNv terminal in a fly expressing ;UAS-mCD8::GFP;Pdf-GAL4 > ;;UAS-sytα::mCherry. The membrane is labeled in green (GFP), and dense core vesicles are labeled in magenta (mCherry). **c**) Magnified views of a single axial frame at a selected region from panel (c), revealing the fine structural details of the vesicles and membrane organization. e) Images of a single varicosity. Left: z-projection. Middle and Right: single frames separated axially by 2 μm.

To further explore the potential of our custom-made LSM, we studied flies expressing GFP in the plasma membrane and mCherry in DCVs.^58^ DCVs are specialized secretory (hence, dynamic) organelles that transport neuropeptides from the soma to the terminals. Figure 4c shows an example of a *z*-projection and Figure 4d shows a close-up of a single selected frame. The 3D data is shown in Supplementary Video 2. The plasma membrane is shown in green and DCVs in magenta. The close-up image shows several DCV puncta decorating *en passant* and terminal varicosities (boutons). Remarkably, the spatial resolution of our method allows the identification of different clusters of DCV inside a single varicosity. Figure 4e shows a single varicosity of approximately 1,6 μm wide. In the z-projection (Figure 4e, left), two clusters of DCVs are visible, laterally separated by 800 nm. In addition, these two clusters are axially offset, with the one on the right located 2 μm above the other (see Figures 4e middle and right). Overall, we have demonstrated conditions for *in vivo* imaging of organelles (or subcellular structures) in a living adult fly with unprecedented spatial resolution.

## Discussion

We designed and built a 2-color upright SOLS microscope for imaging of living adult *Drosophila* flies. Because it utilizes a single objective at the interface with the flies’ head, it offers a series of advantages relative to other LS implementations that require several objectives, such as lattice light sheet microscopy. First, it is compatible with a WF epifluorescence imaging module for simpler and faster sample finding and positioning. Second, it images without mechanically moving the fly or any part in contact with it. Third, and more importantly, it is less invasive because it requires a smaller dissection area on the head’s cuticula, as it is only necessary to accommodate the light cone of a single objective. Furthermore, this configuration also offers open space for other devices, such as perfusion systems. Altogether, these advantages have a positive impact on the flies’ health, which is crucial for long-term imaging experiments.

These features were already available with the most used technology for live imaging of *Drosophila* which is 2pM. However, the SOLS microscopy presented here avoids using high-power infrared pulses that are potentially phototoxic. Moreover, the LS selective illumination guarantees significantly lower phototoxicity and photobleaching. Here, transgenic living flies expressing plasma membrane GFP were imaged 75 times before observing a 50% drop in the emitted fluorescence signal. This number is more than ten times larger than what can be achieved with confocal microscopy, with similar image quality.

While it has been previously demonstrated that SOLS microscopy is capable of achieving resolutions close to the diffraction limit on a series of living animals such as zebrafish or *Drosophila* eggs,^42^ here we have demonstrated it for brain imaging in a living adult fruit fly for the first time. We demonstrate a lateral resolution as high as 370 nm for 488-nm excitation in an axial FOV of 12 μm. Remarkably, this is comparable to the achievable resolutions of 2pM.^59,60^ The axial diffraction-limited FOV can be extended up to 30 μm without significantly losing lateral resolution, but at the expense of axial resolution and sectioning.

We also show two-color imaging on the axonal terminals of sLNv neurons. We imaged ransgenic flies expressing a fluorescent dye at the membrane in combination with either the mitochondria or the dense-core vesicles. This type of measurements could enable studies that correlate neuronal structure with the (dynamic) distribution of specific organelles of interest. This contrasts with most published research on live imaging of *Drosophila*, which typically studies calcium dynamics in response to external stimuli. In particular, long-term *in vivo* imaging of sLNv could provide insights into the circadian plasticity of these clock neurons. Until now, these studies are typically cross-sectional using a different individual for each time point.^4^

Overall, the combined demonstration of reduced invasiveness, phototoxicity and photobleaching, in addition to the high spatial resolution, can pave the way for future long-term experiments in living *Drosophila*.

Importantly, our custom-built microscope allows for easily implementable modifications. These could include higher NA primary objectives for even higher resolution and optical throughput. There is a supportive community of SOLS microscope builders and developers,^61^which facilitates the incorporation of all the recent and expanding advancements of SOLS microscopy, such as the combination of LS with optogenetic stimulation,^41^ multi-view imaging,^42,62^ live-projection imaging,^63^ Raman spectroscopy,^64^ structured illumination,^65^ or adaptive optics.^66^

## Materials and Methods

### Microscope components and specifications

**O1:** 60x, 1.1 NA, LUMFLN60XW, Olympus. **TL1:** SWTLU-C f =180 mm, Olympus. **O2:** 60x, 0.9 NA, air, MPLAPON60X, Olympus. **TL2:** Effective focal length 135 mm - TTL200MP + ACT508-750-A + ACT508-500-A, Thorlabs. **O3:** AMS-AGY-v1.0, Applied Scientific Instrumentation. **TL3:** TTL200-A - f = 200 mm, Thorlabs **SL1 and SL2:** f =70 mm CLS-SL, Thorlabs. **L1:** AC254-030-A-ML - f=30 mm, Thorlabs **CL1:** ACY254-250-A - f = 250.0 mm, Thorlabs **CL2:** ACY254-050-A - f = 50.0 mm, Thorlabs **CL3:** ACY254-100-A - f = 100.0 mm, Thorlabs. **GM:** Saturn 9B, ScannerMAX. **Lasers:** LBX-488 and LCX-561, Oxxius. **Cameras:** pco.edge 4.2 sCMOS, PCO and CS165MU/M - Zelux®, Thorlabs. **BS:** BSW26R 50:50, Thorlabs. **DM1, DM2:** ZT405/488/561rpc-UF3, Chroma. **NF:** ZET488/561m, Chroma. **EF:** ET510/20m for 488 nm excitation and ET630/75m for 561 nm excitation, Chroma. The effective FOV_y_ = 113 μm is given by 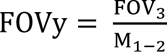, where FOV_3_ = 150 μm and M_1-2_ = 1.33 M1−2 The effective (NA_x_, NA_y_) = (1.1, 1) of the composed optical system was calculated using the *Crossbill Design* software.^67^ The parameters used in the calculations were: MO1=60x, NA1=1.1, *n*_1_=1.33, MO2=60x, NA2=0.9, *n*_2_=1, TL1=180 mm, TL2=135 mm, SL1=70 mm, SL2=70 mm, MO= 40x, NA3= 1, *n*_3_= 1.52, TL3= 200 mm, Tilt angle= 41°, FN1 = 26.5, FN2= 26.5 and FN3= 6.

### Subject details

All experiments were performed on adult flies, 2–5 days post-eclosion. Stocks ;*Pdf*-GAL4;*Pdf*-GAL4 (#80939), used as a driver for s-LNvs and ;UAS-mCD8::GFP; (#5137), used to label plasma membrane, were obtained from Bloomington Stock Center. ;;UAS-Sytα::mCherry, employed to label dense-core vesicles, was generously provided by Paul H. Taghert.^58^ ;UAS-mito::mKO; was generated in our lab to label mitochondria.

### Fly mounting and preparation

An aluminum foil with a hand-made hole was glued to the 3D-printed fly holder (Fig 3a), following published methods.^55^ The fly was anesthetized on ice and under a dissection microscope, its head and thorax were positioned into the hole using forceps (Fine Science Tools). The head of the fly was bent forward to give access to its posterior surface and carefully immobilized by the eyes employing beeswax (70 °C, Sigma) and a thin-end wax carving pencil.

Under saline solution (103 mM NaCl, 3 mM KCl, 5 mM TES, 8 mM trehalose, 10 mM glucose, 26 mM NaHCO3, 1 mM NaH2PO4, 4 mM MgCl2 and 1.5 mM CaCl2),^68^ the cuticle in the dorso-posterior region of the head was removed employing sharpened #5 forceps and a tungsten needle (Fine Science Tools). The air sacs and the fat tissue were also removed to obtain optical access to the s-LNv region. The M16 muscle was cut to prevent brain motion.

### Data acquisition

The microscope’s components are controlled through a Python-based graphical user interface specifically programmed for image acquisition tasks. The acquisition sequence was adapted from the code available at the GitHub repository https://github.com/amsikking/SOLS_microscope. Image acquisition was synchronized via a National Instruments board, which managed the triggering between the camera and the galvo mirror driver. The exposure time for the volumes shown in Figures 3 and 4 was 100 ms per frame. This resulted in a total acquisition time of around 6 seconds for a 60-frame volume. The excitation power measured before the entrance pupil of O1 was 1 mW, giving an irradiance at the focal plane of 30 μWμm^-2^. For the measurements presented in Figures 3e and 3f, a lower irradiances of 15 and 7,5 μWμm^-2^ were used, respectively. Sequential acquisition of two channels was acquired, and channels were subsequently combined computationally using ImageJ.^69^ Images in Figure 3e and all images of Figure 4 went through a Richardson–Lucy deconvolution process with 10 iterations.^70^

### Image Deskewing

Acquired images were deskewed to correct for oblique imaging geometry. The deskewing process was implemented based on the code provided by the QI2lab on GitHub (https://github.com/QI2lab/OPM/tree/master).^41^ This approach corrected the image stacks by realigning the oblique planes, resulting in a corrected 3D volume. Using this code, the angle and step size of the light sheet were accounted for, enabling accurate reconstruction of the imaged neurons. The deskewed images were processed and combined for downstream analysis in ImageJ.

### Beads samples and PSF measurements

Beads samples were made on commercial glass slides. The glass first underwent a cleaning process: it was rinsed with acetone and abundant distilled water, sonicated in a 0.2% Hellmanex solution for ten minutes, and dried at 100°C. Then, the glass was positively charged using poly- (diallyl-dimethylammonium chloride) (PDDA, Sigma-Aldrich, MW ≈ 400–500 kg mol^−1^). They were immersed in a solution of 1 mg/ml PDDA in 0.5 M NaCl. Finally, a drop containing the beads was deposited on the glass for a few minutes and rinsed with water afterwards. The beads solution was 40 nm, 505/515, Invitrogen for the 488 nm laser characterization, and 100 nm, 580/605, Invitrogen for the 561 nm laser characterization.

The fabricated sample was used to characterize the PSF. Although the SOLS microscope acquires volumes, the beads sample has emitters on a single plane. Therefore, by acquiring a volume, only information from the PSF at the sample position is obtained. Figure S2a shows an example of one of the acquired images.

After acquisition and deskewing, only the frame corresponding to the axial position of the sample was analysed, using the software Picasso.^71^ This software localizes the emitters and fits a 2d Gaussian function for each emitter. Figure S1b shows a histogram of the measured values. The PSF in the z-axis was calculated using the PSFj software with the same set of data.^72^

### Decorrelation analysis

To estimate the achieved resolution, a parameter-free estimation based on the decorrelation analysis presented by Descloux was employed.^73^ The analysis was performed over the z-stack presented in Figure 3b. Figure S1 shows the lateral resolution as a function of the depth within the sample. Within the range of 4 μm, the resolution remains approximately constant at 544 nm with a standard deviation of 32 nm.

## Acknowledgments

We thank Lucía Lopez, Gonzalo Escalante, Alan Szalai, and Fernando Stefani for support and materials.

We thank Alfred Millett-Sikking, Kayley Hake, Juan Carlos Boffi, Luciano Masullo and Andrew York for insightful advice and discussions.

We thank Eugenia Chiappe for tips and training on sample preparation.

F.J.T. holds a fellowship from the National Agency for the Promotion of Science, Technology and Innovation (Agencia I+D+i). J.I.I., M.RC are supported by graduate fellowships from the Argentine Research Council for Science and Technology (CONICET); J.G. and M.F.C. are members of CONICET; This work was supported by the PICT2018-0995 (to M.F.C.) from Agencia I+D+i, Argentina and by R01NS108934 (to H. delaI., M.E. and M.F.C.). The funders had no role in study design, data collection and analysis, decision to publish, or preparation of the manuscript.

## Author contributions

Research conception: M.F.C.

Microscope design: M.B. and J.G

Microscope building and alignment: F.J.T. and J.G.

Software development for microscope control: F.J.T.

Data acquisition: F.J.T., L.S, M.F, M.RC, J.I.I. and J.G.

Data analysis: F.J.T., M.B. L.S, M.F, and J.G.

Supervision and Funding: M.E., H.O.I, and M.F.C.

Fly Pushing and dissection: F.J.T., M.RC, and J.I.I.

Writing—original draft: M.B., and J.G.

Figures: F.J.T., M.B, L.S, M.F, and J.G.

Writing—review & editing: All authors.

## Data and materials availability

All data generated or analyzed during this study are included in this published article (and its supplementary information files).

## References

1. Schlegel, P. et al. Whole-brain annotation and multi-connectome cell typing of Drosophila. Nature 634, (2024).

2. Wilson, R. I., Turner, G. C. & Laurent, G. Transformation of Olfactory Representations in the Drosophila Antennal Lobe. Science (80-). 303, 366–370 (2004).

3. Mann, K., Gallen, C. L. & Clandinin, T. R. Whole-Brain Calcium Imaging Reveals an Intrinsic Functional Network in Drosophila. Curr. Biol. 27, 2389–2396.e4 (2017).

4. Huang, C. et al. Long-term optical brain imaging in live adult fruit flies. Nat. Commun. 9, 1–10 (2018).

5. Sinha, S. et al. High-speed laser microsurgery of alert fruit flies for fluorescence imaging of neural activity. Proc. Natl. Acad. Sci. 110, 18374–18379 (2013).

6. Hsu, K.-J., Lin, Y.-Y., Chiang, A.-S. & Chu, S.-W. Optical properties of adult Drosophila brains in one-, two-, and three-photon microscopy. Biomed. Opt. Express 10, 1627 (2019).

7. Chen, Z., Truong, T. M. & Ai, H. W. Illuminating brain activities with fluorescent protein-based biosensors. Chemosensors 5, 1–28 (2017).

8. Górska-Andrzejak, J. et al. Circadian expression of the presynaptic active zone protein bruchpilot in the lamina of Drosophila melanogaster. Dev. Neurobiol. 73, 14–26 (2013).

9. Fernández, M. P., Berni, J. & Ceriani, M. F. Circadian Remodeling of Neuronal Circuits Involved in Rhythmic Behavior. PLOS Biol. 6, e69 (2008).

10. Gorostiza, E. A., Depetris-Chauvin, A., Frenkel, L., Pírez, N. & Ceriani, M. F. Circadian pacemaker neurons change synaptic contacts across the day. Curr. Biol. 24, 2161–2167 (2014).

11. Duhart, J. M. et al. Modulation and neural correlates of postmating sleep plasticity in Drosophila females. Curr. Biol. 33, 2702–2716.e3 (2023).

12. Wang, Q. et al. Optical control of ERK and AKT signaling promotes axon regeneration and functional recovery of PNS and CNS in Drosophila. Elife 9, e57395 (2020).

13. Grienberger, C., Giovannucci, A., Zeiger, W. & Portera-Cailliau, C. Two-photon calcium imaging of neuronal activity. Nat. Rev. Methods Prim. 2, (2022).

14. Sinha, S. et al. Principles of Two-Photon Excitation Microscopy and Its Applications to Neuroscience. 110, 823–839 (2013).

15. Isaacman-Beck, J. et al. SPARC enables genetic manipulation of precise proportions of cells. Nat. Neurosci. 23, 1168–1175 (2020).

16. Wang, J. W., Wong, A. M., Flores, J., Vosshall, L. B. & Axel, R. Two-photon calcium imaging reveals an odor-evoked map of activity in the fly brain. Cell 112, 271–282 (2003).

17. Pacheco, D. A., Thiberge, S. Y., Pnevmatikakis, E. & Murthy, M. Auditory activity is diverse and widespread throughout the central brain of Drosophila. Nat. Neurosci. 24, 93– 104 (2021).

18. Lu, R. et al. Video-rate volumetric functional imaging of the brain at synaptic resolution. Nat. Neurosci. 20, 620–628 (2017).

19. Leong, J. C. S., Esch, J. J., Poole, B., Ganguli, S. & Clandinin, T. R. Direction selectivity in drosophila emerges from preferred-direction enhancement and null-direction suppression. J. Neurosci. 36, 8078–8092 (2016).

20. Seelig, J. D. et al. Two-photon calcium imaging from head-fixed Drosophila during optomotor walking behavior. Nat. Methods 7, 535–540 (2010).

21. Brezovec, B. E. et al. Mapping the neural dynamics of locomotion across the Drosophila brain. Curr. Biol. 34, 710–726.e4 (2024).

22. Tainton-Heap, L. A. L. et al. A Paradoxical Kind of Sleep in Drosophila melanogaster. Curr. Biol. 31, 578–590.e6 (2021).

23. Münch, D., Goldschmidt, D. & Ribeiro, C. The neuronal logic of how internal states control food choice. Nature vol. 607 (Springer US, 2022).

24. Mann, K., Deny, S., Ganguli, S. & Clandinin, T. R. Coupling of activity, metabolism and behaviour across the Drosophila brain. Nature 593, 244–248 (2021).

25. Picot, A. et al. Temperature Rise under Two-Photon Optogenetic Brain Stimulation. Cell Rep. 24, 1243–1253.e5 (2018).

26. Donovan, E. J. et al. Dendrite architecture determines mitochondrial distribution patterns in vivo. Cell Rep. 43, 114190 (2024).

27. Delestro, F. et al. In vivo large-scale analysis of Drosophila neuronal calcium traces by automated tracking of single somata. Sci. Rep. 10, 1–14 (2020).

28. Harris, D. T., Kallman, B. R., Mullaney, B. C. & Scott, K. Representations of Taste Modality in the Drosophila Brain. Neuron 86, 1449–1460 (2015).

29. Aimon, S., Cheng, K. Y., Gjorgjieva, J. & Kadow, I. C. G. Global change in brain state during spontaneous and forced walk in drosophila is composed of combined activity patterns of different neuron classes. eLife vol. 12 (2023).

30. Aimon, S., et al. Fast near-whole-brain imaging in adult drosophila during responses to stimuli and behavior. PLoS Biology vol. 17 (2019).

31. Lu, Z. et al. Long-term intravital subcellular imaging with confocal scanning light-field microscopy. Nat. Biotechnol. (2024).

32. Hillman, E. M. C., Voleti, V., Li, W. & Yu, H. Light-Sheet Microscopy in Neuroscience. Annu. Rev. Neurosci. 42, 295–313 (2019).

33. Stelzer, E. H. K. et al. Light sheet fluorescence microscopy. Nat. Rev. Methods Prim. 1, (2021).

34. Lemon, W. C. et al. Whole-central nervous system functional imaging in larval Drosophila. Nat. Commun. 6, (2015).

35. Ahrens, M. B., Orger, M. B., Robson, D. N., Li, J. M. & Keller, P. J. Whole-brain functional imaging at cellular resolution using light-sheet microscopy. Nat. Methods 10, 413–420 (2013).

36. Liang, X., Holy, T. E. & Taghert, P. H. Synchronous Drosophila circadian pacemakers display nonsynchronous Ca2+ rhythms in vivo. Science (80-). 351, 976–981 (2016).

37. Hubert, A. et al. Enhanced neuroimaging with a calcium sensor in ex-vivo Drosophila melanogaster brains using closed-loop adaptive optics light-sheet fluorescence microscopy. J. Biomed. Opt. 28, (2023).

38. Strack, R. Single-objective light sheet microscopy. Nat. Methods 18, 28 (2021).

39. Dunsby, C. Optically sectioned imaging by oblique plane microscopy. Opt. InfoBase Conf. Pap. 16, 186–196 (2009).

40. Kumar, M., Kishore, S., Nasenbeny, J., McLean, D. L. & Kozorovitskiy, Y. Integrated one- and two-photon scanned oblique plane illumination (SOPi) microscopy for rapid volumetric imaging. Opt. Express 26, 13027 (2018).

41. Sapoznik, E. et al. A versatile oblique plane microscope for large-scale and high-resolution imaging of subcellular dynamics. Elife 9, 1–39 (2020).

42. Yang, B. et al. DaXi—high-resolution, large imaging volume and multi-view single-objective light-sheet microscopy. Nat. Methods 19, 461–469 (2022).

43. Yang, B. et al. Epi-illumination SPIM for volumetric imaging with high spatial-temporal resolution. Nat. Methods 16, 501–504 (2019).

44. Bouchard, M. B. et al. Swept confocally-aligned planar excitation (SCAPE) microscopy for high-speed volumetric imaging of behaving organisms. Nat. Photonics 9, 113–119 (2015).

45. Voleti, V. et al. Real-time volumetric microscopy of in vivo dynamics and large-scale samples with SCAPE 2.0. Nat. Methods 16, 1054–1062 (2019).

46. Schaffer, E. S. et al. The spatial and temporal structure of neural activity across the fly brain. Nat. Commun. 14, (2023).

47. Botcherby, E. J., Juškaitis, R., Booth, M. J. & Wilson, T. An optical technique for remote focusing in microscopy. Opt. Commun. 281, 880–887 (2008).

48. Millett-Sikking, A., Thayer, N. H., Bohnert, A. & York, A. G. calico/remote\_refocus: Pre-print. (2018) doi:10.5281/zenodo.1146084.

49. Millett-Sikking, A. & York, A. AndrewGYork/high_na_single_objective_lightsheet: Work-in-progress. (2019) doi:10.5281/ZENODO.3376243.

50. Kumar, M. & Kozorovitskiy, Y. Tilt (in)variant lateral scan in oblique plane microscopy: a geometrical optics approach. Biomed. Opt. Express 11, 3346 (2020).

51. Lvarez, P. A. L. O. Z. A. Light-sheet microscopy : a tutorial. 10, 111–179 (2018).

52. Stoleru, D., Peng, Y., Nawathean, P. & Rosbash, M. A resetting signal between Drosophila pacemakers synchronizes morning and evening activity. Nature 438, 238–242 (2005).

53. Renn, S. C. P., Park, J. H., Rosbash, M., Hall, J. C. & Taghert, P. H. A pdf neuropeptide gene mutation and ablation of PDF neurons each cause severe abnormalities of behavioral circadian rhythms in Drosophila. Cell 101, 791–802 (2000).

54. Helfrich-Förster, C. & Homberg, U. Pigment-dispersing hormone-immunoreactive neurons in the nervous system of wild-type Drosophila melanogaster and of several mutants with altered circadian rhythmicity. J. Comp. Neurol. 337, 177–190 (1993).

55. Jayaraman, V. & Laurent, G. Evaluating a genetically encoded optical sensor of neural activity using electrophysiology in intact adult fruit flies. Front. Neural Circuits 1, 1–9 (2007).

56. Ji, I. et al. Ultrastructural correlates of circadian structural plasticity. 22, (2024).

57. Jolivet, N. & Bertolin, G. Revealing mitochondrial architecture and functions with single molecule localization microscopy. Biol. Cell 117, 1–19 (2025).

58. Park, D., Li, P., Dani, A. & Taghert, P. H. Peptidergic Cell-Specific Synaptotagmins in *Drosophila*: Localization to Dense-Core Granules and Regulation by the bHLH Protein DIMMED. J. Neurosci. 34, 13195 LP – 13207 (2014).

59. Benninger, R. K. P. & Piston, D. W. Two-photon excitation microscopy for the study of living cells and tissues. Current Protocols in Cell Biology (2013).

60. Peres, C., Nardin, C., Yang, G. & Mammano, F. Commercially derived versatile optical architecture for two-photon STED, wavelength mixing and label-free microscopy. Biomed. Opt. Express 13, 1410–1429 (2022).

61. Millett-Sikking, A. Snoutclub. (2025) doi:10.5281/zenodo.14861847.

62. Sparks, H., Almagro, J., Behrens, A., Salbreux, G. & Dunsby, C. Dual-view oblique plane microscopy. Opt. InfoBase Conf. Pap. 11, 7204–7220 (2021).

63. Chen, B. et al. Projective light-sheet microscopy with flexible parameter selection. Nat. Commun. 15, 1–8 (2024).

64. Guo, K., Kalyviotis, K., Pantazis, P. & Rowlands, C. J. Hyperspectral oblique plane microscopy enables spontaneous, label-free imaging of biological dynamic processes in live animals. Proc. Natl. Acad. Sci. U. S. A. 121, 1–9 (2024).

65. Chen, B. et al. Resolution doubling in light-sheet microscopy via oblique plane structured illumination. Nat. Methods 19, 1419–1426 (2022).

66. Mcfadden, C. et al. Adaptive optics in an oblique plane microscope. Biomed. Opt. Express 15, 4498 (2024).

67. Kumar, M. & KozorovitskiyLab. KozorovitskiyLaboratory/Crossbill-Design: Crossbill Design. (2019) doi:10.5281/zenodo.3554784.

68. Fujiwara, T., Brotas, M. & Chiappe, M. E. Walking strides direct rapid and flexible recruitment of visual circuits for course control in Drosophila. Neuron 110, 2124–2138.e8 (2022).

69. Rueden, C. T. et al. ImageJ2: ImageJ for the next generation of scientific image data. BMC Bioinformatics 18, 1–26 (2017).

70. Richardson, W. H. Bayesian-Based Iterative Method of Image Restoration*. J. Opt. Soc. Am. 62, 55–59 (1972).

71. Schnitzbauer, J., Strauss, M. T., Schlichthaerle, T., Schueder, F. & Jungmann, R. Super-resolution microscopy with DNA-PAINT. Nat. Protoc. 12, 1198–1228 (2017).

72. Theer, P., Mongis, C. & Knop, M. PSFj: know your fluorescence microscope. Nat. Methods 11, 981–982 (2014).

73. Descloux, A., Grußmayer, K. S. & Radenovic, A. Parameter-free image resolution estimation based on decorrelation analysis. Nat. Methods 16, 918–924 (2019).

