## Supplementary Information for "Single Objective Light Sheet Microscopy allows high-resolution *in vivo* brain imaging of Drosophila"

### Supplementary Note 1. Measurement of the beam profile

The light sheet XZ profile was measured by scanning a single 100 nm fluorescent bead through the excitation sheet of light at 561 nm. For sample preparation, a dilution 1:$10^7$ from stock (580/605, Invitrogen) is drop casted for 1 minute on top of a PMMA-coated clean glass slide and then rinsed thoroughly with distilled water.

The bead was scanned through the light sheet using a motorized stage with a step size of 0.5 µm. At each sample position, an image was acquired with the LS camera. The total detected fluorescence intensity at each bead position was calculated by integrating the total counts on a region of interest of 60x60 px. The resulting normalized intensity profile is shown in Figure 2c, and used to calculate the following beam parameters:

1. Beam Waist: The transverse intensity profile at the beam waist (x' axis) was fitted with a Gaussian function, as shown in Figure 2d. The full width at half-maximum (FWHM) of this curve represents the beam waist, $w_0$= (2.2 ± 0.1) µm
2. Light Sheet Length (Lz'): Figure S1 shows a linear profile along the *z' axis*, corresponding to the dashed line from Figure 2c. The light sheet length $L_z$ = (39 ± 1) µm was computed as twice the distance over which the intensity drops by a factor of $\sqrt{2}$.

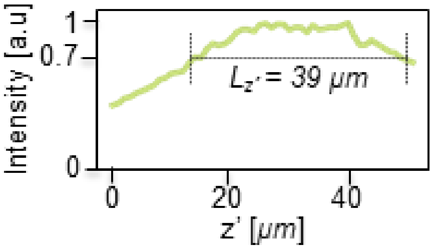


***Figure S1)*** *Normalized intensity profiles of the LS in the z' axis.*

For the measurement of the Light Sheet Width (Ly), a glass sample containing a higher concentration of beads was prepared. The LS module was used to obtain a 3D image of the sample. The frame corresponding to the XY plane containing the beads is shown in Figure S2a. Figure S2b shows an intensity profile along the y axis, corresponding to the dashed line of Figure S2a. The resulting FWHM is $L_y$=(53 ± 1) µm.

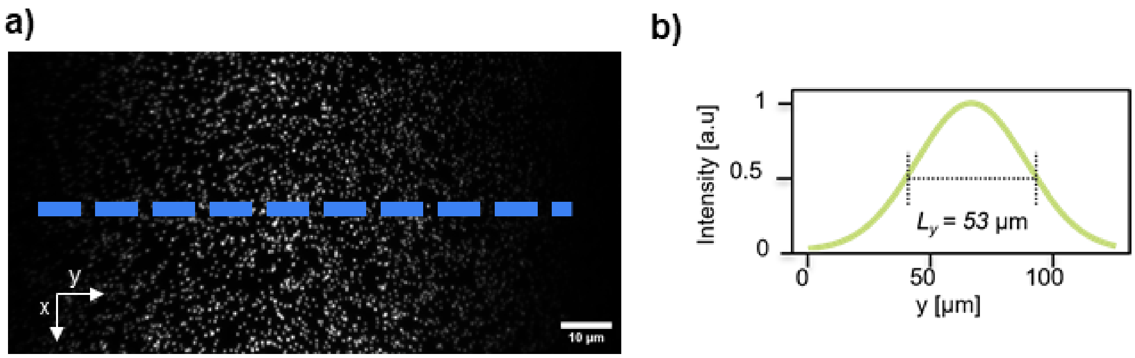


***Figure S2 a)*** *Normalized intensity profiles of the LS in y axis* ***b)*** *Normalized intensity profiles of the LS in the z' axis.*

<u>Supplementary Note 2. Measurement of the Point Spread Function using fluorescent beads.</u>

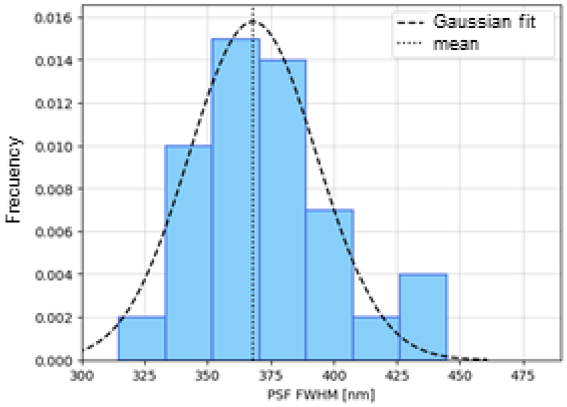


***Figure S3)*** *Histogram of the measured FWHM for the PSF at the objective focal plane, under 488 nm excitation.*

<u>Supplementary Note 3. Parameter-free resolution estimation based on decorrelation analysis</u>

Figure S3 shows the lateral resolution as a function of the depth within the sample. We used *the z-stack shown in Figure 3b (main text).* Within the range of 4 µm, the resolution remains approximately constant, with a mean value of 544 nm and a standard deviation of 32 nm.

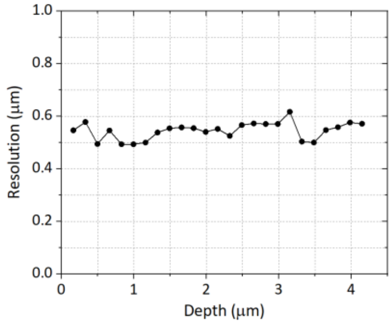


***Figure S3)*** *Lateral resolution as a function of depth.*

### Supplementary Note 4. Photobleaching comparison between Confocal and SOLS

In this section, we compare the SOLS microscope with a Zeiss LSM 710 confocal microscope equipped with a W Plan-Apochromat 20×/1.0 objective. The comparison between two distinct imaging techniques is not straightforward, due to the inherent differences in illumination and detection geometry, excitation irradiance, exposure time, detector, etc. In addition, the photobleaching rate depends on the acquisition parameters. For these reasons, we performed a comparison between techniques at comparable image quality. We used the signal to noise ratio (SNR) as a metric of the image quality. In both microscopes, neuronal terminals of the small lateral ventral neurons (s-LNvs) were imaged, using live adult flies with the same genotype ;UAS-mCD8::GFP;*Pdf*-GAL4, excited at 488 nm, with the same wavelength detection window (500 - 520) nm. The imaging conditions are summarized in Table S1.

| Microscope | I $[\frac{nW}{(nm)^2}]$ | T [seg] | SNR | SBR | Volumes |
|---|---|---|---|---|---|
| Confocal | 7 | 76 | 22 ± 4 | 32 ± 2 | (6 ± 2) |
| SOLS | 4 x $10^{-3}$ | 1 | 26 ± 4 | 6 ± 3 | (75 ± 10) |

***Table 1)*** *Comparison of photobleaching experiments. It includes the maximum irradiance (I) at the sample, acquisition time required to image 80×50×12 $\mu m^3$ (T), signal-to-noise ratio (SNR), signal-to-background ratio (SBR), and the number of volumes acquired before the signal dropped to half.*

Figures S4a and S4b show an example of a frame corresponding to the initial acquired volume, for confocal and SOLS, respectively. The calculation of the SNR of each image was as follows. First, we identified small features in the images, small enough to be diffraction-limited and follow a Gaussian shape, as the ones marked with red lines in Figures S4a and S4b. Then, we plotted the intensity profile, fitted a Gaussian function defined as $f(x) = ae^{-\frac{(x-b)^2}{2c^2}} + d$, and calculated the root mean square deviation (RMSD) of the fit. Finally, the SNR is computed as SNR = $\frac{a}{RMSD}$ and the signal to background ratio SBR = $\frac{a}{d}$. Figures S4c and S4d shows the intensity profiles (in black) corresponding to the red lines of Figures S4a and S4b, for the initial frame of confocal and SOLS microscopy, respectively. The Gaussian fits are shown in blue.

For the selected imaging parameters, the confocal microscope has a SNR = (22 ± 4), while SOLS had (26 ± 4). On the other hand, the confocal microscope had a SBR = (32 ± 2), while SOLS had (6 ± 3).

For each imaging session in the confocal microscope, 10 volumetric acquisitions were performed, with the intensity dropping to half after (6 ± 2) acquisitions. In the SOLS, 200 volumes were acquired, with the intensity dropping to half after (75 ± 10) acquisitions.

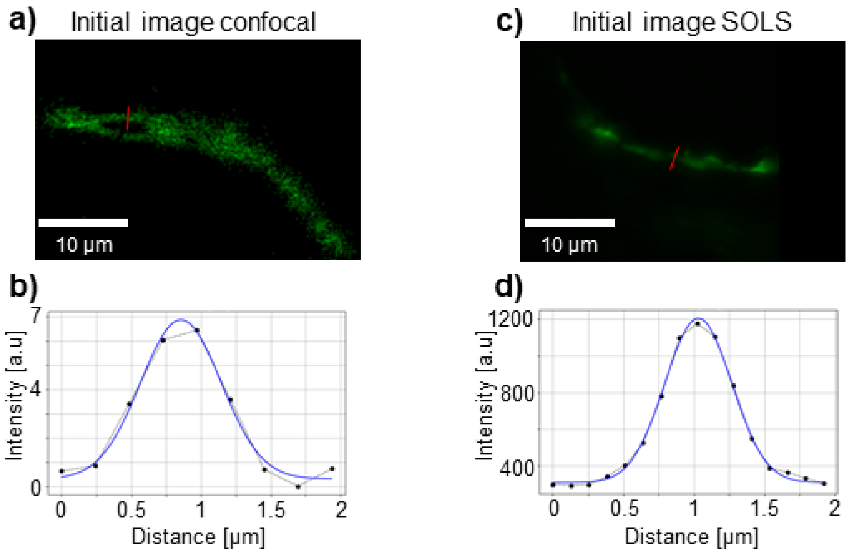


***Figure S4 a)*** *A frame from the initial volume acquired with the confocal microscope. The red line indicates the region where the Gaussian profile was fitted.* ***b)*** *The intensity profile extracted from the confocal image, with raw data shown in black and the Gaussian fit in blue.* ***c)*** *A frame from the initial volume acquired with the SOLS microscope. The red line indicates the region where the Gaussian profile was fitted.* ***d)*** *The intensity profile extracted from the SOLS image, with raw data in black and the Gaussian fit in blue.*
